# Expanding Clinical Phage Microbiology: Simulating Phage Inhalation for Respiratory Tract Infections

**DOI:** 10.1101/2021.06.14.448272

**Authors:** Shira Ben Porat, Daniel Gelman, Ortal Yerushalmy, Sivan Alkalay-Oren, Shunit Coppenhagen-Glazer, Malena Cohen-Cymberknoh, Eitan Kerem, Israel Amirav, Ran Nir-Paz, Ronen Hazan

**Affiliations:** Institute of Biomedical and Oral Research (IBOR), Faculty of Dental Medicine, The Hebrew University of Jerusalem, Jerusalem, Israel; Department of Military Medicine, Faculty of Medicine, The Hebrew University of Jerusalem, Jerusalem, Israel; Department of Clinical Microbiology and Infectious Diseases, Hadassah-Hebrew University Medical Center, Jerusalem, Israel; Pediatric Pulmonology Unit and Cystic fibrosis Center, Hadassah Medical Center; Faculty of Medicine, Hebrew University of Jerusalem, Israel; Pediatric Pulmonary Unit, Dana-Dwek Children’s Hospital, Tel Aviv

## Abstract

Phage therapy is a promising antibacterial strategy for resistant respiratory tract infections. Phage inhalation may serve this goal; however, it requires a careful assessment of their delivery by this approach. Here we present an *in-vitro* model to evaluate phage inhalation.

Eight phages, most of which target CF-common pathogens, were aerosolized and administered to a real-scale CT□derived 3D airways model with a breathing simulator. Viable phage loads reaching the output of the nebulizer and the tracheal level of the model were determined and compared to the loaded amount.

Phage inhalation resulted in a diverse range of titer reduction, primarily associated with the nebulization process. No correlation was found between phage delivery to the phage physical or genomic dimensions. These findings highlight the need for tailored simulations of phage delivery, ideally by a patient-specific model in addition to proper phage matching, to increase the potential of phage therapy success.

**Take-Home Message:** Phage therapy can be used against infectious diseases if personally tailored. Using a 3D airways model, we show that phage delivery by inhalation to the respiratory tract is unpredictable and also requires a precise evaluation.

## Introduction

Phage therapy refers to the use of bacteriophages (phages), bacterial viruses, as antimicrobial agents. Lytic phages can propagate in the presence of their bacterial hosts while sequentially lysing proximal bacterial cells. This enables them to penetrate and destroy biofilm and co-evolve with bacterial targets [1]. The emerging threat of antimicrobial resistance has re-introduced this previously neglected method and its advantages to the clinical practice, with an increasing number of reports applying phages for infectious diseases in recent years [2–4].

A significant target of phage therapy is the treatment of life-threatening pulmonary infections [5]. In this regard, phages have been used in cases of cystic fibrosis (CF), an inherited life-shortening disease associated with recurrent and chronic lung infections [6–8]. Examples include the use of intravenous (IV) phages in a 26-year-old CF patient with (MDR) *Pseudomonas aeruginosa* pneumonia [7], a case of a 12-year-old lung-transplanted CF patient treated by inhaled phages and by direct phage administration via therapeutic bronchoscopy for *Achromobacter xylosoxidans* [8], and the treatment of six individuals with CF or non-CF bronchiectasis, treated by nebulized phages for *P. aeruginosa* [9]. Evidently, these published cases differ in treatment protocols and specifically in the method for phage delivery.

The current understanding of the importance of a personalized and accurate phage matching for therapy dictates the use of simulations and *in-vitro* predictions during phage therapy treatment design [10]. Phage inhalation is not exceptional in this manner, as differences in the biochemical and physical characteristics of phages may cause sub-optimal delivery to the target site and thus should be tested in advance.

Several *in-vitro* studies have previously compared phage recovery at the output of various nebulizers [11, 12]. However, phage concentration reaching the lower respiratory tract was usually not measured but only estimated, for instance, according to the aerodynamic diameter of the released particles [11]. To bridge the gap, we applied a real-scale 3D model of patient’s CT□derived airways. This model, which was previously used for *in-vitro* aerosol studies [13, 14] was adapted here for evaluation of phage delivery to the lungs by inhalation.

## Methods

### Phages and bacteria

Phages targeting the following seven bacterial strains, most of which are associated with CF, were used (Table 1): *P. aeruginosa* strains PA14 and PAR1, *Mycobacterium abscessus* MAC107, *Burkholderia cepacia* strains BCC129 and BCC378, *Staphylococcus aureus* SAR1 and *Staphylococcus epidermidis* SE52. Except for the laboratory strain *P. aeruginosa* PA14, all isolates were obtained from the Clinical Microbiology Department at the Hadassah Medical Center, Israel. *S. epidermidis* was grown in brain heart infusion (BHI) broth (Difco, Sparks, MD) and all other bacteria were grown in LB broth (Difco, Sparks, MD), all at 37 ° C under aerobic conditions shaken on a rotary shaker at 0.85 rcf. BHI or LB agar (1.5%) plates were respectively used for isolation streaks and phage enumeration. All phages were isolated as part of the Israeli Phage Bank [15]. The phages were propagated on their respective target bacterial hosts as previously described [16].

**Table 1.** Characteristics of the phages used in the study.

| Phage | Host | Taxonomy | Genome size<br>(bp) | Genome<br>Structure | GenBank accession |
| --- | --- | --- | --- | --- | --- |
| **BCSR5** | *B. cepacia* BCC378 | Myoviridae | 227,351 | Circular | MW460245.1 |
| **SAOMS1** | *S. aureus* SAR1 | Herelleviridae | 140,135 | Circular | MW460250.1 |
| **BCSR129** | *B. cepacia* BCC197 | Myoviridae | 66,147 | Circular | MW460247.1 |
| **PASA16** | *P. aeruginosa* PA14 | Myoviridae | 66,127 | Circular | MT933737.1c |
| **Itty13** | *P. aeruginosa* PAR1 | Podoviridae | 61,818 | Linear | MW460249.1 |
| **Maco2** | *M. abscessus* c107 | Siphoviridae | 48,239 | Circular | MW460248.1 |
| **Maco7** | *M. abscessus* c107 | Siphoviridae | 47,836 | Linear | MZ152914 |
| **SeAlphi** | *S. epidermidis* SE52 | Podoviridae | 18,368 | Circular | MZ152915 |

### Phage enumeration

Phage titers were determined by the double-layered agarose assay as previously described [17]. Briefly, agar plates were covered with pre-warmed agarose (0.5%), to which 300 μl of the relevant overnight bacterial culture was added. The phage lysates were serially diluted by 10-fold, and 5 μl drops of these dilutions were spotted on the double-layered plates which were then incubated overnight at 37 °C. The number of plaques on the plates was then counted to calculate the initial phage titers, presented as plaque-forming units (PFU)/ml.

### Phage recovery from filters

Phages were recovered from filters placed across the model to evaluate phage viability at these sites. Following each run of the nebulizer, the used filter was left overnight in a sterile tube containing 15 ml of fresh media in 4° C, followed by phage enumeration.

To estimate the fraction of phage titer reduction caused by the used filters (*Fr* -filter reduction fraction), 0.2 ml of each phage in a known concentration were dropped directly on the pads. The filters were then left to dry, and later treated as describe above. For each phage, the *Fr* was calculated as the proportion between viable phages retrieved from the filters to their original inserted titer (Table 2).

**Table 2.**
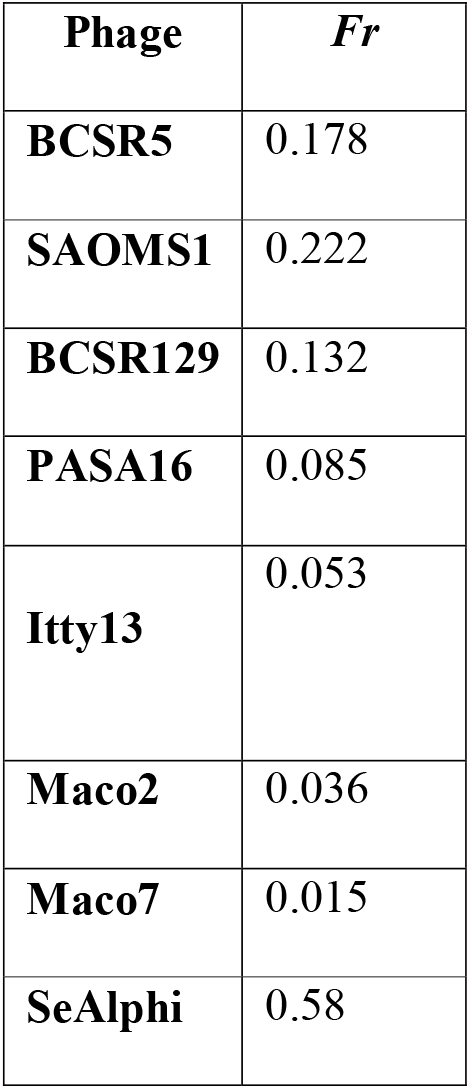
Filter reduction fraction (Fr) of each phage.

### Phage nebulization

The commonly clinically used Pari LC Sprint nebulizer (Pari GmbH, Starnberg, Germany), combined with Pari TurboBoy SX Compressor (Pari GmbH, Starnberg, Germany) was used for phage nebulization (Figure 1). In each run, 3 ml of phage lysate in a previously determined concentration were pipetted into the inhalation device reservoir, and the compressor was operated for 15 minutes. For evaluation of phage viability following nebulization, a PARI filter/valve set (Pari GmbH, Starnberg, Germany) filled with single-use filter pads (Pari GmbH, Starnberg, Germany) was connected to the output of the nebulizer to collect aerosol particles (Figure 1A).

**Figure 1.**
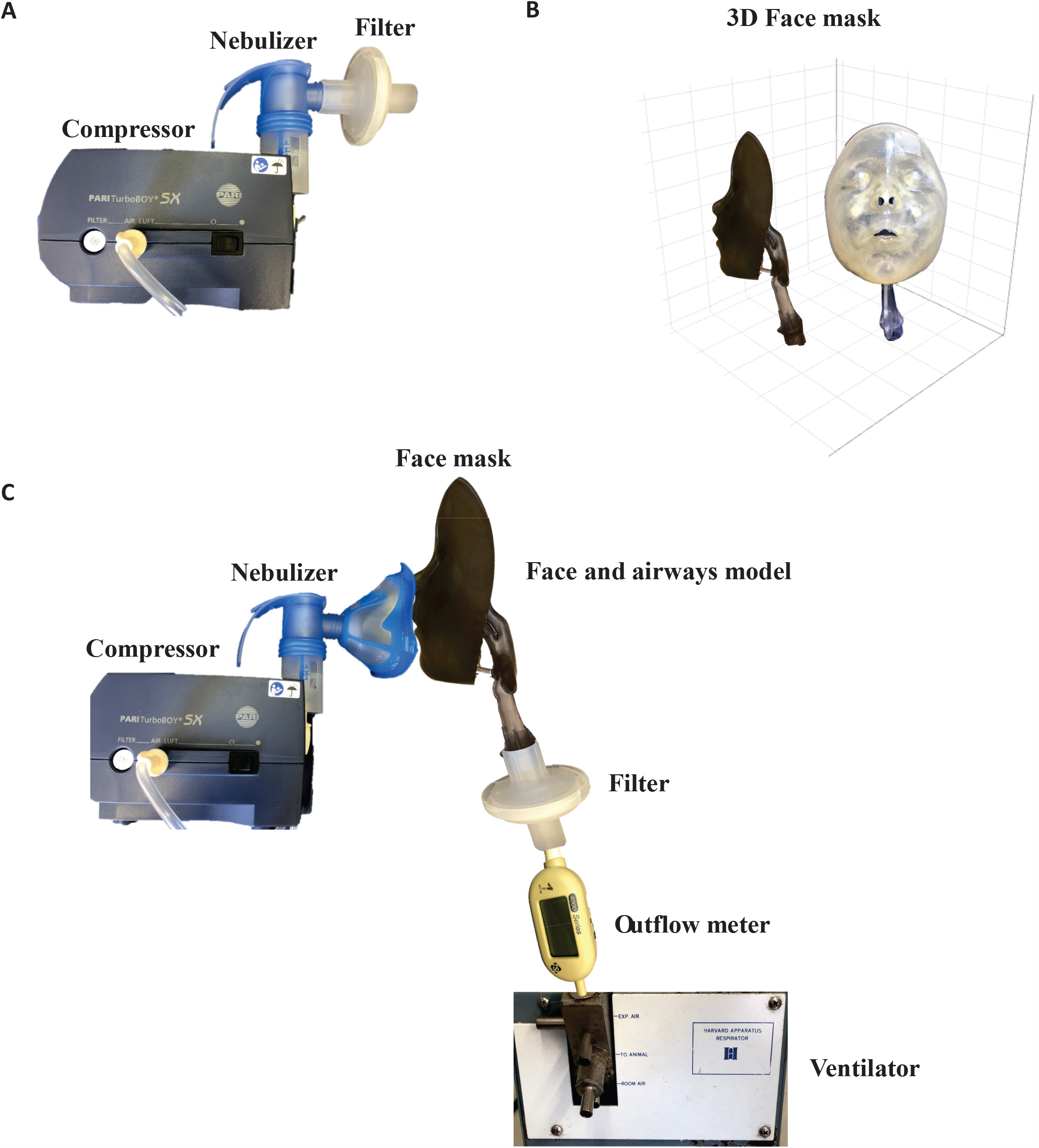
Model for evaluation of phage nebulization and delivery. (A) Demonstration and schematic representation of phage nebulization. Filter pads were placed at the output of the nebulizer to allow phage capture after each run of nebulization. (B) Anterior and lateral views of the 3D face and respiratory airways model. (C) Demonstration and schematic representation of phage inhalation. Phages were nebulized and delivered by a face mask to the face and respiratory tract model, connected to a breathing simulator. Phages were captured by filter pads placed at the distal orifice of the airways model.

### 3D face and airways model construction

The ability of nebulized phages to reach the lower respiratory tract (lung delivery) was evaluated using a 3D reconstruction model of a face and respiratory tract airways [13, 14] (Figure 1B). For the face and airways model creation, CT (Brilliance CT, 64-channel scanner, Philips Healthcare, Best, The Netherlands) scans of 5-year-old children with no craniofacial anomalies were obtained for medical indications. The data were digitized and converted to electronic files. Technical acquisition parameters included: 64 x 0.625 collimation, 0.75-s pitch, 120 kV, 100 mA, and a 2-s rotation time. The thickness of a slice was 2.5 mm, with increments of 1.25 mm. Slices were stored in Digital Imaging and Communications in Medicine (DICOM) format. Subsequently, the slices were joined together to compose a 3-dimensional reconstruction image, which was stored in Standard Transformation Language format, and later transferred over local area network (LAN) for further analysis. The scans were reconstructed and stored in stereolithography (STL) format from which the model was then printed using rapid prototype development techniques. We constructed the model using a photopolymer resin (PolyJet FullCure 720, Objet Geometries Ltd., Sint-Stevens-Woluwe, Belgium), a transparent, stiff material widely used in rapid product development techniques on an Objet Eden 330 3D Printer (Objet Geometries, Rehovot, Israel). Printing layer thickness was 0.016 mm. Anatomical structures included were all air conducting parts from the nostrils to 5 mm below the glottis.

In addition, a breathing simulator (Harvard Apparatus Respirator Model 665, Harvard Apparatus, USA) was connected through the distal orifice of the model. The breathing simulator generated a standard waveform at a pre□set frequency (f=20□breaths/min) and tidal volume (Vt=80□ml), representing breathing patterns appropriate for the model size. A flow meter (TSI Model 4043, TSI Inc., USA) was connected to the breathing simulator and to the filter/valve set to ensure the flow characteristics through the model (Figure 1C).

### Lung delivery of phages

The face and airways model was connected to the Pari LC Sprint nebulizer (Pari GmbH, Starnberg, Germany) by a tightly connected face mask (Pari GmbH, Starnberg, Germany) at the output of the nebulizer (Figure 1C). In each run, 3 ml of phage lysate were pipetted into the nebulizer reservoir, and the compressor was operated for 15 minutes. For evaluation of phage viability at the tracheal level, the PARI filter/valve set (Pari GmbH, Starnberg, Germany) was now placed at the distal portion of the airways model to collect aerosol particles reaching this site (Figure 1C).

### Phage physical characteristics determination

Transmission electron microscopy (TEM) was used for measurement of phage particles size. Phage samples were prepared as previously described [16]. Briefly, 1 ml of phage lysate containing at least 10^8^ PFU/ml was centrifuged at 12,225 g (WiseSpin® CF□10, Daihan Scientific) for 1.5 h at room temperature. The supernatant was discarded, and the pellet was resuspended in 200 μl of 5 mM MgSO4 and left for 24 hours at 4°C. For grid preparation, 10 μl of the phage mixtures were added to 30 μl of 5 mM MgSO4, and the grids were placed on the drops with the carbon side facing down. After a minute, the grid was placed on a 30 μl drop of NanoVan (Nanoprobes, NY, USA), followed by an additional incubation for 5-10 sec. A transmission electron microscope (Jeol, TEM 1400 plus) with a charge-coupled device camera (Gatan Orius 600) was used to capture images. Phage particle’s size was determined as an average of measurements capturing 4-14 (median 8.5) virions for each phage, using ImageJ 1.53h software (http://imagej.nih.gov/ij/).

### Statistical analysis

Phage titers were compared by an unpaired *t-test*. A significance level of 0.05 was used to determine the statistical difference of viable phage loads due to nebulization or lung delivery. Pearson’s correlation test was used for correlation plots. GraphPad prism 8.02 Build 263 (GraphPad Software, Inc, USA) was used for statistical tests. Adobe Illustrator (CC release 21.0.2) was used for graph drawing.

## Results

### The effect of inhalation on viable phage titers

We evaluated the delivery of eight different phages to the lungs using an anatomically accurate face and airways model with corresponding simulation of breathing patterns. The tested phages were selected due to their ability to target pathogens isolated from CF patients (Table 1). An additional phage targeting *S. epidermidis* was tested due to a remarkable difference in morphology.

The phages were nebulized and administered to the model. After phage titer correction according to the *Fr* (Table 2), we found a diverse range of titer drops during phage inhalation to the lungs between the various phages (Figure 2). This reduction ranged from 2.8-log for the phage Maco7 to only 0.2-log reduction with the phage SeAlphi. Overall, seven of the eight tested phages presented a significant titer drop at the tracheal level, simulating their deposition in the lower respiratory tract (Figure 2).

**Figure 2.**
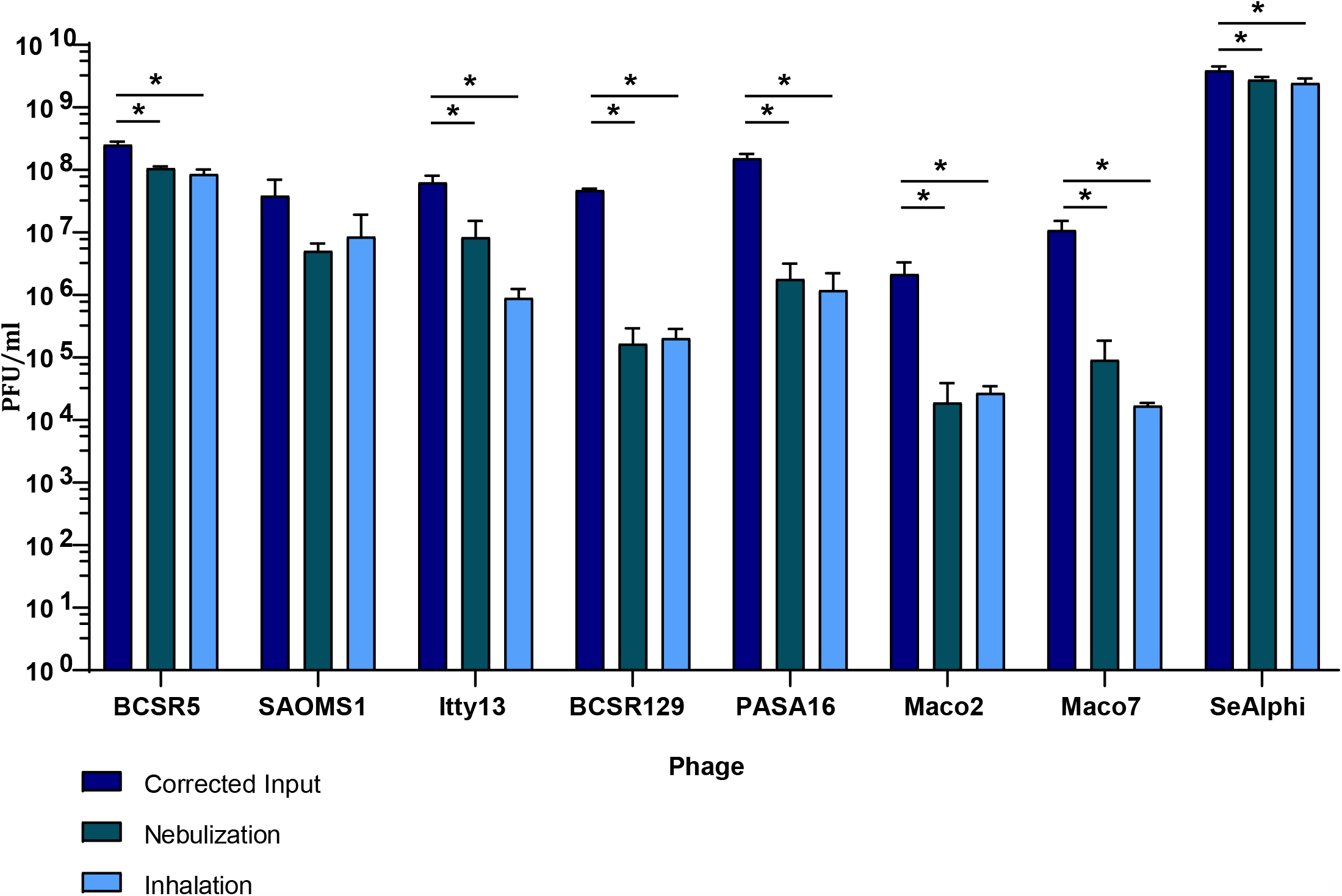
Titer reduction of phages following nebulization and inhalation to the airways model. For each phage, the initial phage titer inserted to the nebulizer, corrected according to the filter reduction fraction (Corrected Input), is compared to the phage titers retrieved from filters placed at the output of the nebulizer (Nebulization) and at the tracheal level of the respiratory tract model (Inhalation). Three separate runs were made for each phage in every condition. Asterisks represent significant difference (p<0.05).

### Determination of phage titer drop during the process of nebulization

In order to better understand the factors associated with phage titer reduction during inhalation, we evaluated phage viability following the nebulization process alone. Viable aerosolized phages were now captured at the output of the nebulizer, representing their titer at the oropharyngeal level. We found that a significant reduction in viable phage loads was also observed following nebulization for all phages presenting decreased titers at the tracheal level (Figure 2). The nebulization process also led to a diverse range of phage titer reductions, between 2.45-log for the phage BCSR129 to only 0.15-log for the phage SeAlphi. Furthermore, in all phages presenting a significant titer drop at the tracheal level, no significant difference was found between their loads at the trachea and at the output of the nebulizer. This demonstrates that this reduction is highly due to the nebulization process itself.

### Effect of phage characteristics on titer drop during inhalation

Next, we turned to test for intrinsic phage characteristics which may affect their delivery. That way, if a correlation between specific phage characteristics and their ability to pass the model is found, it will allow to better predict the desired phages to administer and their required dosage. The tested traits included the size of the capsids, tail length, total phage length and genome size. However, no correlation was found between these tested characteristics and the phage deposition at the tracheal level (Figure 3).

**Figure 3.**
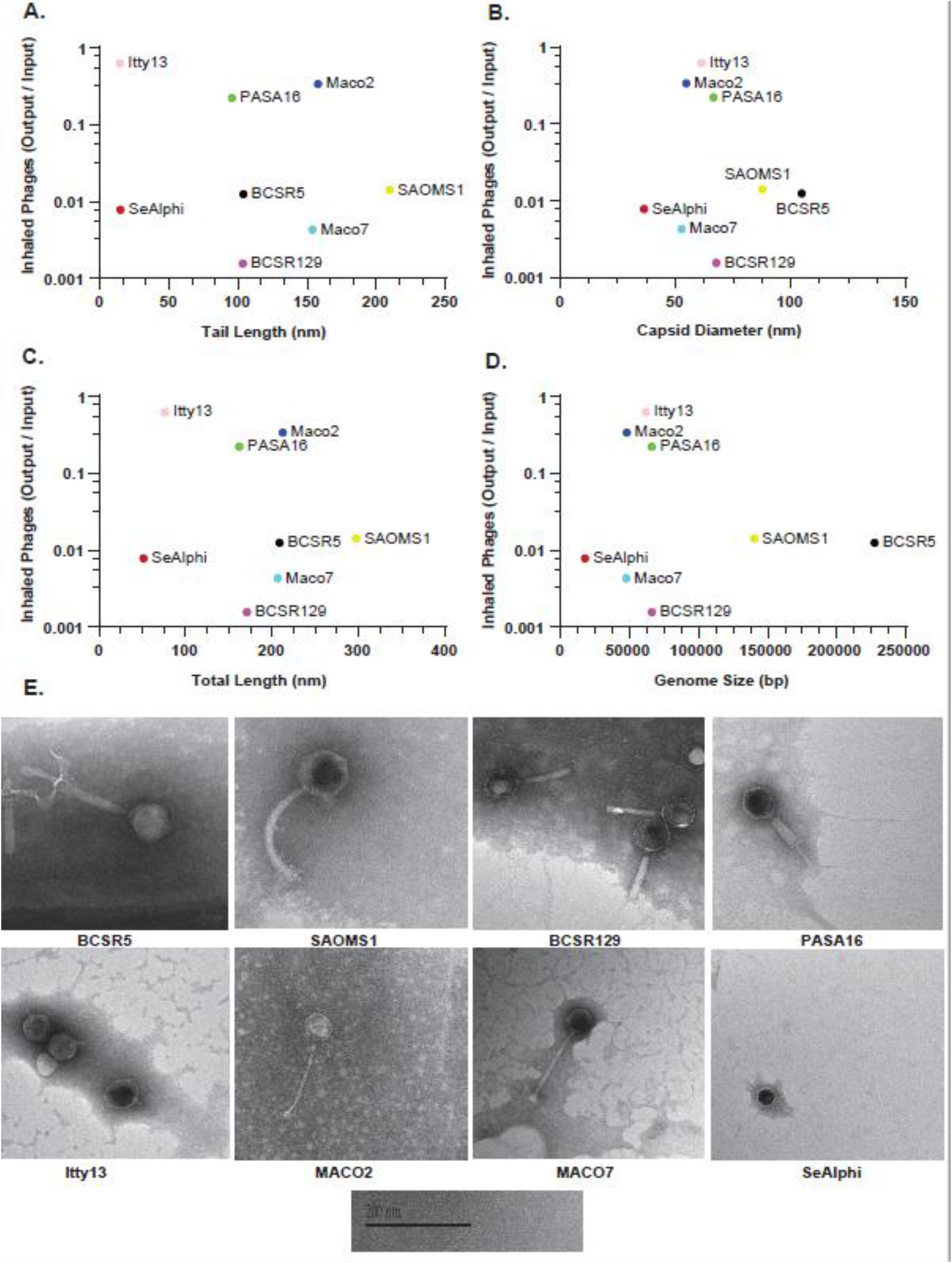
Phage characterization and its effect on inhalation fitness. (A-D) Correlation plots representing phage inhalation efficacy, defined as the proportion between viable phages reaching the tracheal level and the corrected initial phage loads, according to (A) tail length, (B) capsid diameter, (C) total phage length and (D) phage genome size. (E) TEM images of the phages used in the study. All images are scaled according to the bar representing 200 nm.

## Discussion

In this study we have simulated phage inhalation using a dedicated real size 3D reconstruction model of human airways, with an adjusted breathing simulator. To the best of our knowledge, this is a first of a kind simulation allowing to directly measure phage delivery to the lungs in realistic human aerodynamic characteristics. Using this model, we found that there is a high variation in the levels of phage titer reduction following inhalation among different phages. Moreover, this reduction was mostly attributed to the nebulization process itself and was significant in seven of eight tested phages.

Previous reports have evaluated the ability of phages to be aerosolized, usually by direct measurements of their titers at the output of the nebulization device. Sahota *et al*. described the recovery of specific anti-Pseudomonal phages following nebulization [11]. In their study, the total number of viable phages leaving a jet nebulizer ranged between 15% for one phage to 2% for another respectively to their original amount. Carrigy *et al*. have shown that the same anti-Tuberculosis phage presented highly diverse levels of titer reductions, ranging between 0.4-and 3.7-logs, when nebulized by different devices such as a vibrating mesh nebulizer, a jet nebulizer and a soft mist inhaler [12]. In another study, Astudillo *et al*. have supplied visual proof for the structural changes of phages upon nebulization by different methodologies [18]. Using TEM, they visualized that nebulization of a specific anti-Pseudomonal phage by jet, vibrating-mesh or static-mesh nebulizers resulted in high levels of phage tail separation. As in the current study, these reports emphasize that phage titer may drop during nebulization, and that the most appropriate phages chosen for inhalation, as well as the nebulization methodology itself may vary.

Leung *et al*. have suggested that nebulization-induced titer loss was particularly correlated with the tail length of the phage [19]. In their study, viable phage titers dropped by 0.04-to 2-logs following jet nebulization, in correlation with phage morphology. However, in our study, using different phages with a wide range of different characteristics, including short-tailed Podoviridae as well as Myoviridae and Siphoviridae, no correlation to tail length was observed. Furthermore, we have also found no correlation between phage inhalation capabilities to phage size, genome size, taxonomy or to the bacterial host. Thus, we conclude that routine genomic and morphological characterization of the phages may not be sufficient to predict their nebulization and inhalation abilities. Accordingly, specific *in-vitro* evaluations should be performed to address phage delivery prior to inhalation treatment. This point was recently demonstrated by Guillon *et al*., who measured phage delivery by nebulization *in-vitro*, to select the most appropriate inhalation interface and phage dosage before administering aerosolized phages to pigs suffering from pneumonia [20].

The current study has several limitations. Firstly, although the presented model supplies accurate anatomical characteristics, it does not include the complete physiological conditions found in the respiratory tract. These include the complex tissue environment, the mucocilliary structures and the immune system effects. Thus, an additional reduction in phage titers may be expected in the clinical setting. Accordingly, the evaluation of these parameters on phage inhalation requires additional studies using animal models, which will further expand the knowledge in the field [20, 21]. Secondly, even though many phages from diverse families were included in the study, a wider evaluation could potentially shed more light on specific characteristics associated with titer decrease during inhalation.

Finally, we have used only one nebulizer type, while other nebulizers may behave differently [12]. However, this only reinforces our main conclusion, that prior testing of the phage and the inhalation device in a tailored model are important for proper treatment design.

In conclusion, here we have shown that as in many other cases, phages may act in a non-predictive way in terms of lung deposition following inhalation. Thus, analogously to the concept of phage matching which is based on accurate laboratory testing of phages against the bacterial target [10], *in-vitro* phage delivery assays should also be performed in a personalized manner, potentially by patient-specific airways models in the future. In this regard, proper phage inhalation requires not only the selection of the most suitable phage, but also the most appropriate inhalation device by empirical *in-vitro* assessments.

## Acknowledgements

The authors thank Yossi Aldar for technical assistance, and Ori Inbar from the Cystic Fibrosis foundation in Israel for insightful ideas.

